# Integrated Multi-Omics Analysis of Brain Aging in Female Nonhuman Primates Reveals Altered Signaling Pathways Relevant to Age-Related Disorders

**DOI:** 10.1101/2022.11.01.514742

**Authors:** Laura A. Cox, Sobha Puppala, Jeannie Chan, Kip D. Zimmerman, Zeeshan Hamid, Isaac Ampong, Hillary F. Huber, Ge Li, Avinash Y. L. Jadhav, Benlian Wang, Cun Li, Mark G. Baxter, Carol Shively, Geoffrey D. Clarke, Thomas C. Register, Peter W. Nathanielsz, Michael Olivier

## Abstract

The prefrontal cortex (PFC) has been implicated as a key brain region responsible for age-related cognitive decline. Little is known about aging-related molecular changes in PFC that may mediate these effects. To date, no studies have used untargeted discovery methods with integrated analyses to determine PFC molecular changes in healthy female primates. We quantified PFC changes associated with healthy aging in female baboons by integrating multiple omics data types (transcriptomics, proteomics, metabolomics) from samples across the adult age span. Our integrated omics approach using unbiased weighted gene co-expression network analysis (WGCNA) to integrate data and treat age as a continuous variable, revealed highly interconnected known and novel pathways associated with PFC aging. We found GABA tissue content associated with these signaling pathways, providing one potential biomarker to assess PFC changes with age. These highly coordinated pathway changes during aging may represent early steps for aging-related decline in PFC functions, such as learning and memory, and provide potential biomarkers to assess cognitive status in humans.

## 1 Introduction

The prefrontal cortex (PFC) is central to working memory, temporal processing, decision making, flexibility, and goal-oriented behavior (Funahashi and Andreau 2013; Friedman and Robbins 2022). Studies in humans and multiple nonhuman primate (NHP) species have shown reductions of PFC activity and changes in neuronal morphology without loss of neurons in aging that are associated with cognitive decline (Lacreuse et al., 2020; Upright and Baxter 2021). Ianov *et al*., (Ianov et al., 2016) observed age-related differences in gene expression in the PFC of young (5–6 months, *n* = 11) and aged (17–22 months, *n* = 20) male Fischer 344 rats; however, these were not associated with age-related cognitive impairment in a PFC-mediated task. Erraji-Benchekroun *et al*., (Erraji-Benchekroun et al., 2005) observed age-related transcriptional changes in Broadmann’s Area 9 (BA9) and BA47 in 39 humans from 13 to 79 years of age; however, half of these were samples obtained after suicides and the postmortem interval (PMI) averaged 17.5 hours, both of which very likely effect RNA and protein quality.

Previous studies on primate PFC changes with age, which include humans and NHPs, focused on gene and miRNA expression (PMID: 22162950, PMID: 22783978, PMID: 20591156), RNA editing (PMID: 24152549), RNA splicing (PMID: 23340839), and associations between mRNA and their encoded proteins (PMID: 25853883). However, these studies included only males, with all but one study including less than 7 across the adult age span. A few of these studies included one elderly female – consequently little is known about aging-related PFC changes in female primates. To date, there is no detailed dataset available characterizing multi-omic molecular changes in the PFC across the adult age span in healthy primates.

The goal of this study was to integrate data from multiple omics methods to quantify PFC molecular changes associated with healthy aging in female baboons (*Papio*), a NHP model of aging. This study is unique compared to prior studies in that it evaluates a relatively large group of NHPs (n=34) across the entire adult age span (human equivalent ∼30 to 88 years). All animals consumed the same healthy diet, were group-housed the same way, and tissues were collected on a defined schedule with short PMIs (∼30 min); therefore, this type of controlled investigation is not possible in humans. In addition, previous studies used RNA-Seq data (PMID: 22162950, PMID: 22783978, PMID: 20591156) (PMID: 24152549) and mRNA associated proteins (PMID: 25853883) to quantify molecular changes associated with age; whereas, our study integrated transcriptomic, proteomic, and metabolomic data. In previous studies, we (Cox et al., 2021) and others (e.g., (Meng et al., 2019) have demonstrated greater power to detect phenotypically relevant molecular pathways using integrated omics compared to transcriptomics analyses alone.

Our integrated omics approach using weighted gene co-expression network analysis (WGCNA), an unbiased method to construct molecular networks based on pairwise correlations between variables, and pathway enrichment analysis revealed 2 modules containing 587 transcripts and 13 proteins negatively correlated with age. We identified an additional 57 proteins and 20 metabolites associated with age using regression analyses. Pathway enrichment analysis revealed 25 overlapping, coordinated pathways negatively correlated with age. The top ranking pathways included dopamine-DARPP32 feedback in cAMP signaling and estrogen receptor signaling, both previously associated with age-related changes in PFC (Luine and Frankfurt 2013; Bailey et al., 2011; Arnsten et al., 1994). In addition, top ranking pathways included pathways not previously associated with aging-related PFC changes - synaptogenesis, synaptic long term depression, and nitric oxide signaling. We also identified proteins negatively correlated with age that serve as key regulators in these signaling pathways, and age-associated metabolites that are regulators or by-products of these pathways. Our untargeted integrated omic approach revealed highly coordinated, novel pathways that may represent early events leading to aging-related decline in PFC functions, such as learning and memory, and provide potential biomarkers to assess cognitive status in humans.

## 2. Materials and Methods

### 2.1 Animal Care and Maintenance

The study included 34 females ranging in age from 7.5y to 22.1y (human equivalent ∼30y to 88y), median age 14.3y (human equivalent ∼50y) (Bronikowski et al., 2002). All procedures were approved by the Texas Biomedical Research Institute (TBRI) Animal Care and Use Committee and conducted in facilities approved by the Association for Assessment and Accreditation of Laboratory Animal Care. The TBRI animal use programs operated according to all National Institutes of Health (NIH) and U.S. Department of Agriculture guidelines, and were directed by board certified veterinarians (DVM). All animal care decisions were made by the Southwest National Primate Research Center (SNPRC) veterinarians.

Baboons (*Papio hamadryas* spp., crosses of olive, hamadryas, and yellow baboons) were housed in outdoor social groups of 3-19 animals at the SNPRC at TBRI, in San Antonio, Texas. Animals were fed monkey chow (Monkey Diet 5LEO, Purina, St Louis, MO, USA) *ad libitum* throughout life, and water was continuously available with multiple lixits in each enclosure. All animals were provided complete veterinary care by SNPRC veterinary staff throughout their lives.

### 2.2 Necropsy

Baboons were pre-medicated with ketamine hydrochloride (10 mg/kg IM) and anesthetized using isoflurane (2%) as previously described (Schlabritz-Loutsevitch et al., 2007). All collections were conducted between 8:00 AM – 10:00 AM to minimize variation from circadian rhythms. While under general anesthesia, baboons were exsanguinated as approved by the American Veterinary Medical Association. Following cardiac asystole, 3mm deep samples of gray matter from the frontal cortex on the left side of the brain, extending from the posterior part of the precentral sulcus to the intersection of the precentral sulcus and lateral sulcus, to the anterior frontal brain were collected (Broca’s area 8,9,10, 44, 45 and 46). Tissues were snap frozen in liquid nitrogen, and stored at -80°C.

### 2.3 Morphometric Measures

Morphometric measures were collected from sedated animals prior to necropsy using standard anatomical landmarks as described previously (Chavez et al., 2009).

### 2.4 Clinical Measures

Blood samples were collected from the femoral vein in overnight fasted animals after intramuscular administration of ketamine at 10 mg/kg. All collections were conducted between 8:00 AM – 10:00 AM to minimize variation from circadian rhythms. Blood samples were collected within 5 min of ketamine administration. For all measures, assay precision was determined by testing pooled samples using 5 replicates in each assay. These assays were repeated at 2 dilutions to assess linearity of the results. All test samples were run at dilutions estimated to achieve values in the middle of the assay calibration range.

Plasma Lipids and Glucose - Total plasma cholesterol (TPC), low density lipoprotein (LDL) cholesterol, high density lipoprotein (HDL) cholesterol, triglyceride, and glucose concentrations were determined by the Wake Forest Comparative Medicine Clinical Chemistry and Endocrinology Laboratory using reagents (ACE) and instrumentation (ACE AXCEL autoanalyzer) from Alfa Wasserman Diagnostic Technologies (West Caldwell, NJ). Plasma lipids were standardized to calibrated controls from the Centers for Disease Control and Prevention/National Institutes of Health Lipid Standardization Program (Solomon Park, Burien, WA, USA).

Leptin - Leptin concentrations were determined by radioimmunoassay using a kit from EMD Millipore (Burlington, MA, kit HL-81K).

### 2.5 Transcriptomics

#### 2.5.1 RNA Isolation

Approximately 5 mg of frozen prefrontal cortex was homogenized in 1 ml RLT buffer (Qiagen) using a BeadBeater (BioSpec) with zirconia/silica beads, and RNA was extracted using the Zymo Direct-zol RNA Miniprep Plus kit according to manufacturer’s instructions. RNA quality was assessed using high sensitivity RNA ScreenTape with the TapeStation instrument (Agilent). RNA was stored at −80°C.

#### 2.5.2 Sequence Data Generation

The Kapa RNA HyperPrep kit with RiboErase (Roche) was used to generate cDNA libraries, and quality assessed using Agilent D1000 ScreenTape according to the manufacturer’s protocol. cDNA libraries were pooled and sequenced using v1.5 reagent kit (Illumina) for paired-end sequencing (2×150) on a NovaSeq 6000 Sequencer.

#### 2.5.3 Sequence Data Analysis

Low quality bases with Phred scores below 30 were removed prior to alignment. Trimmed reads were aligned against the olive baboon reference (Panu_3.0, GCF_000264685.3) using HISAT2 (Kim et al., 2015). Aligned reads were quantified using an expectation-maximization algorithm (Xing et al., 2006) with Panu_3.0 annotation (NCBI release 103). The criteria to be counted as paired-end reads were 100% overlap with transcript sequences and skipped regions of junction reads matched the introns of transcripts. Transcripts without read counts across all samples were filtered out, and normalized by the trimmed mean of M values method. Raw read counts were filtered to remove those with less than 30 counts across all samples and further filtered to include only transcripts with 3 or more counts in 50% of the samples, resulting in 25,381 transcripts that passed quality filters.

### 2.6 Proteomics

#### 2.6.1 Sample Processing

Proteomic samples were prepared as described (Hamid et al., 2022). Briefly, approximately 5 mg of each tissue sample was homogenized in Tris buffer, precipitated overnight in acetone at − 20°C, and centrifuged at 12,000 g for 10 min. The protein pellet was dried, reconstituted in 100 mM of ammonium bicarbonate, and quantified. One hundred mg of protein were reduced using dithiothreitol for 1 h at 56°C, alkylated using iodoacetamide for 30 min in dark, and digested overnight with trypsin. Samples were cleaned and desalted using Thermo Scientific Pierce C18 Tips, dried, and reconstituted in 0.1% formic acid.

#### 2.6.2 LC/MS Data Acquisition and Analysis

LC/MS data were acquired as described (Hamid et al., 2022). One mg of each sample was loaded on a PepMap RSLC C18 easy-spray column (3um, 100A, 75um x 15cm) using Easy-nLC 1200 coupled to an Orbitrap Lumos Tribrid Mass Spectrometer (Thermo Scientific), and peptides were separated using a 3 hr gradient of Mobile phase A (0.1% Formic acid in 95:5 Water:Acetonitrile) and Mobile Phase B (0.1% Formic acid in 80:20 Acetonitrile:Water). Peptides were eluted according to the gradient program: 2% to 30% B in 140 min, 30% to 95% B in 30 min and 95% to 100% B in 10 min. Mass spectrometer data were acquired in MS1 scan mode (m/z=375-1800) with a resolution of 120,000 with Automatic Gain Control of 4.0×105 and maximum injection time of 50 ms. MS/MS data acquisition was done using HCD mode at a resolution of 30,000 with an Automatic Gain Control target of 4.0 × 10^5^ and maximum injection time of 50 ms. All data acquisition was done using Thermo Scientific Xcalibur software.

MS raw data were analyzed using MetaMorpheus (Hamid et al., 2022; Miller et al., 2019) using the *P. anubis* reference proteome database from Uniprot with 44,721 entries (UP000028761). Data files were calibrated using the following settings: precursor mass tolerance of 15 ppm, product mass tolerance of 25 ppm with Carbamidomethyl as fixed modification, and oxidation of methionine as variable modification. Trypsin was selected as protease with 2 maximum mixed cleavages and the calibrated data files were converted to mzML format. Post calibration data was searched using G-PTM-D task for incorporation of common biological, metal or artifact PTMs into the search database. A final search was done using the augmented search database with incorporated G-PTM-D based modifications at precursor and product mass tolerance of 5 and 20 ppm respectively. Peptide and protein quantification were done using the FlashLFQ approach and the Match between runs option was enabled. Protein intensities were normalized using global intensity normalization. In the final normalized data missing values were imputed using Random forest imputation workflow (Hamid et al., 2022; Stekhoven and Buhlmann 2012).

### 2.7 Metabolomics

#### 2.7.1 Sample Processing

Extraction of metabolites from brain samples was performed following a protocol adopted from a previously described study (Misra et al., 2019). Briefly, aliquots (15 μL) of brain homogenates were subjected to sequential solvent extraction, once each with 1 mL of acetonitrile:isopropanol:water (3:3:2) and 500 μL of acetonitrile:water (1:1) mixtures at 4°C (Fiehn et al., 2008). Adonitol (2 μL from 10 mg/ml stock) was added to each aliquot prior to the extraction as internal standard. The extracts were then dried under vacuum at 4°C prior to chemical derivatization (silylation reactions). Tubes without samples (blanks) were treated similarly as sample tubes to account for background noise and other sources of contamination. Samples and blanks were sequentially derivatized with methoxyamine hydrochloride (MeOX) and 1% TMCS (2,2,2-Trifluoro-N-methyl-N-(trimethylsilyl)-acetamide, Chlorotrimethylsilane) in N-methyl-N-trimethylsilyl-trifluoroacetamide (MSTFA) or 1% TMCS containing N-(t-butyldimethylsilyl)-N-methyltrifluoroacetamide (MTBSTFA) as described (Misra et al., 2019). This involved addition of 20 μL of MeOX (20 mg/mL) in pyridine to the dry extracts and incubation at 55°C for 1 h followed by addition of 80 μL MTBSTFA and incubation at 60°C for 1 h.

#### 2.7.2 GC/MS Data Acquisition and Analysis

Data were generated with a high-resolution (HR) Orbitrap Mass Spectrometer (Q Exactive Orbitrap MS, Thermo Fisher) coupled to gas chromatography (GC). In all cases, 1 µL of derivatized sample was injected into the TRACE 1310 GC (Thermo Scientific, Austin, TX) in a splitless (SSL) mode at 220°C. Helium was used as a carrier gas and the flow rate was set to 1 mL/min for separation on a Thermo Scientific Trace GOLD TG-5SIL-MS (30 m length × 0.25 mm i.d. × 0.25 μm film thickness) column with an initial oven temperature of 50°C for 0.5 min, followed by an initial gradient of 20°C/min ramp rate.

The final temperature of 300°C was held for 10 min. All eluting peaks were transferred through an auxiliary transfer line into a Q Exactive-GC-MS (Thermo Scientific, Bremen, Germany). The total run time was 25 min. Data were generated in an electron ionization (EI) mode at the standard 70 eV energy, emission current of 50 μA, and an ion source temperature of 250°C. A filament delay of 5.7 min was set to prevent excess reagents from being ionized. High resolution EI spectra were acquired at 60,000 resolution (fwhm at m/z 200) with a mass range of 50 - 650 m/z. The transfer line was set to 230°C. Data acquisition and instrument control were carried out using Xcalibur 4.3 and TraceFinder 4.1 software (Thermo Scientific). Capillary voltage was 3500V with a scan rate of 1 scan/s. Finally, raw data (.raw files) obtained from data acquired by GC/MS were converted to .mzML formats using the open source ProteoWizard’s msConvert software prior to data preprocessing with MS-DIAL 4.6 software (Riken, Japan, and Fiehn Lab, UC Davis, Davis, CA, USA). The MS-DIAL 4.6 open software was used for raw peak extraction and data baseline filtering and baseline calibration, peak alignment, deconvolution analysis, peak annotation, and peak height integration as described (Tsugawa et al., 2015). Key parameters included peak width of 20 scan, a minimum peak height of 10,000 amplitudes was applied for peak detection, sigma window value of 0.5, and EI spectra cutoff of 50,000 amplitudes for deconvolution. For annotation settings, the retention time tolerance was 0.5 min, the m/z tolerance was 0.5 Da, the EI similarity cutoff was 60%, and the annotation score cutoff was 60%. In the alignment parameters setting process, the retention time tolerance was 0.5 min, and retention time factor was 0.5. Spectral library matching for metabolite identification was performed using an in-house and public library consisting of pool EI spectra from MassBank, GNPS, RIKEN, MoNA. Data were further normalized by QC-based-loess normalization prior to log10 transformation and missing values were imputed based on random forest imputation method (Ampong et al., 2022; Dunn et al., 2011; Stekhoven and Buhlmann 2012).

### 2.8 Statistical Analysis of Integrated Omics Data

#### 2.8.1. Weighted Gene Co-expression Network Analysis (WGCNA)

WGCNA was performed with the WGCNA package (Langfelder and Horvath 2008) in R software according to the R package WGCNA protocol (https://horvath.genetics.ucla.edu/html/CoexpressionNetwork/Rpackages/WGCNA/). We used sample clustering to identify potential outliers and found one sample with significantly greater low abundance RNA Seq transcript values, which was removed from all three omics datasets (Langfelder and Horvath 2008). Subsequent analyses included 33 female baboons. A total of 26,622 omics molecules were included in WGCNA. Brain omics data from female baboons were used to generate a correlation-matrix for all pair-wise omics data. Next, the threshold function was used to obtain soft threshold (power-12) to construct an adjacency matrix in accordance with a scale-free network (Zhang and Horvath 2005). This adjacency matrix was then transformed into a topological overlap matrix (TOM) to measure relative gene interconnectedness and proximity. The TOM was then used to calculate the corresponding dissimilarity (1 – TOM). Average linkage hierarchical clustering coupled with the TOM-based dissimilarity was used to group correlated omics data into modules (Zhang and Horvath 2005). More specifically, modules were generated from the Dynamic Tree Cut method for Branch Cutting. The major parameters were set with a deep-split value of 2 to branch splitting and a minimum size cutoff of 50 (minimum cluster size = 50) to avoid abnormal modules in the dendrogram; highly similar modules were merged together with a height cutoff of 0.25. Modules were considered significant if the correlation was ≥ 0.30 and p-value < 0.05. In the resulting network, as neighbors in a cluster shared high topological overlap, the resulting modules likely indicated a common functional class. WGCNA has the advantage of allowing analysis of continuous traits without binning the data for arbitrary phenotypic cutoffs in an analysis because binning data into categories typically translates into loss of power.

#### 2.8.2 Construction of Module-Trait Relationships

The omics modules summarize the main patterns of variation. The first principal component represents the summary of each module and is referred to as the module eigengene (Langfelder and Horvath 2007).

The relationship between module eigengenes and clinical traits was assessed by Pearson correlation; if p-value < 0.05, then the module and clinical trait were regarded as significant. The modules and clinical traits that showed significant (p-value < 0.05) and high correlations <u>(></u> 0.30) were selected for further investigation. A heat map was used for visualization of the correlations of each module-trait relationship.

#### 2.8.3 Proteomics and Metabolomics Correlation with Age

The central question of our study was whether modules of brain omics data correlated with age. The identified modules negatively correlated with age contained few proteins and no metabolites. WGCNA is designed to alleviate the multiple testing burden in transcriptomic data, which is less of an issue for proteomic and metabolomic data due to their smaller numbers of molecules. Consequently, we used Pearson correlation to nominate proteins and metabolites that correlated with age and age-squared.

Although we did not adjust for multiple testing, we addressed this within pathway enrichment analysis, i.e., pathway statistical values for enrichment plus EOP stringent filtering (Nijland et al., 2007).

### 2.9 Pathway and Network Enrichment Analysis

To assess directionality of pathways significantly enriched with molecules from WGCNA and regression analyses, negatively correlated molecules were converted to negative fold change and positive correlations to positive fold change. All molecules from significant modules were imported to Ingenuity Pathway Analysis (IPA) software (Qiagen) for core analysis where pathways were analyzed for significant enrichment of module genes. Right-tailed Fisher’s exact test was used to calculate associations between molecules in the dataset and molecules in annotated pathways, and pathways were ranked by -log p-value (Spradling et al., 2013). A p-value of < 0.01 was considered significant. We used an EOP approach to identify pathways in which activity was biologically consistent between the beginning and end of the pathway - we considered that a pathway is biologically relevant if fold changes of the molecules at both ends of the pathway were consistent with the overall pathway change, i.e., activated or inhibited (Nijland et al., 2007).

## 3. Results

### 3.1 Morphometric and Clinical Measures

Our study included PFC samples and corresponding blood samples from 34 female baboons (*Papio*) ranging in age from 7.5y to 22.1y (human equivalent ∼30y to 88y). PFC samples were rapidly collected in a controlled setting from scheduled necropsies providing high quality tissues for multi-omic analyses.

Morphometry and blood for clinical chemistries were collected immediately prior to necropsy (Supplemental Table 1). Regression analysis showed body length associated with age (p-value = 0.024) and age-squared (p-value < 0.001), and BMI was nominally associated with age (p-value = 0.063).

**Table 1.**
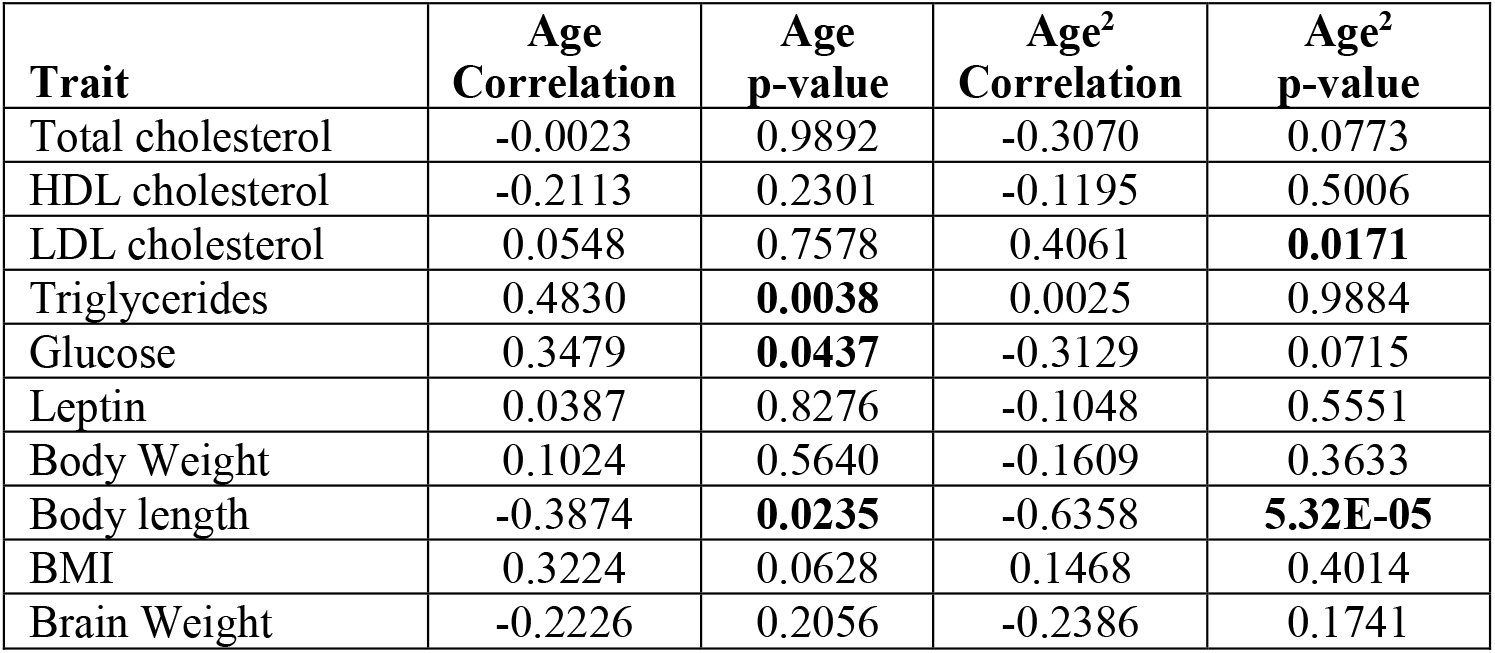
Correlations between age and morphometric and clinical measures

Among the clinical measures, triglycerides (p-value = 0.004) and glucose (p-value = 0.044) were associated with age, and LDL cholesterol was associated with age-squared (p-value = 0.019) (Table 1).

### 3.2 Integrated Omics to Identify Age-Associated Molecules

We identified 25,381 transcripts (Supplemental Table 2), 917 proteins (Supplemental Table 3), and 324 metabolites (Supplemental Table 4) that passed quality filters (Table 2). We used WGCNA to identify 26 modules of co-correlated omics molecules, and then determined whether any of these modules associated with age, age-squared, and morphometric and clinical measures.

**Table 2.** Summary of Genes, Proteins and Metabolites Associated with Age

| Molecule | Quality | WGCNA<br>Age | WGCNA<br>Age<sup>2</sup> | Corr.<br>Age | Corr.<br>Age<sup>2</sup> |
| --- | --- | --- | --- | --- | --- |
| Genes | 25,381 | 587 | 0 | - | - |
| Proteins | 917 | 13 | 0 | 27 | 30 |
| Metabolites | 324 | 0 | 0 | 0 | 20 |

**Table 3.**
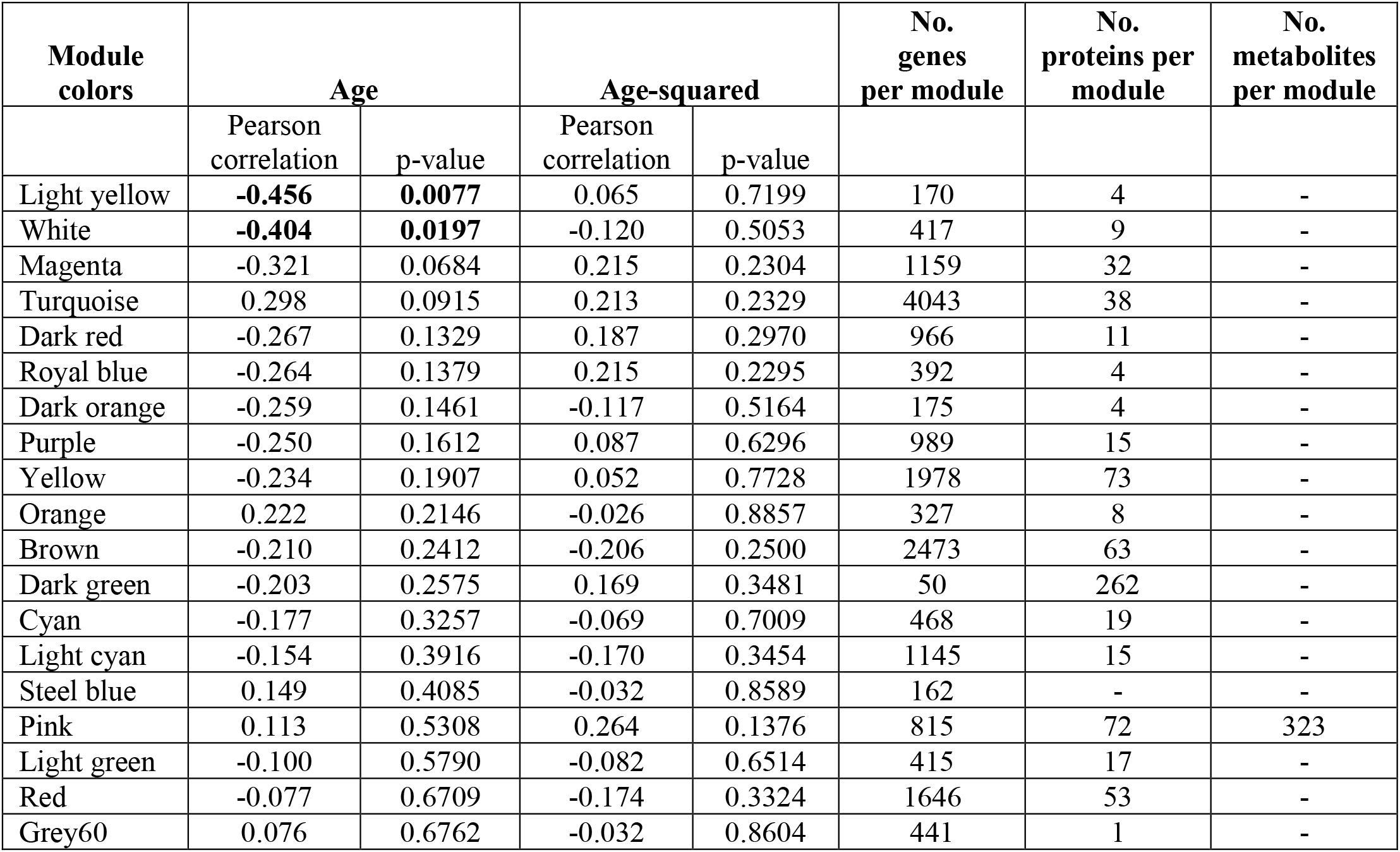
Number of genes, proteins, and metabolites in the top 20 WGCNA modules with statistical results for age and age^2^

We identified 2 modules negatively correlated with age (white, p-value = 0.008, correlation = -0.45; and light yellow, p-value = 0.020, correlation = -0.40) (Table 2, Supplemental Table 5) which included 587 genes, 13 proteins, and 0 metabolites (Supplemental Table 6, Table 1). We did not find any significant modules associated with age-squared. However, we did identify 27 proteins (23 mappable in IPA) negatively correlated with age, 30 proteins associated with age-squared (21 mappable in IPA), and 20 metabolites associated with age-squared (Supplemental Table 7).

To determine whether metabolic and/or morphometric measures were associated with PFC molecular changes with age, we assessed whether age-associated WGCNA modules overlapped with morphometric- and clinical measures-associated WGCNA modules. Although triglycerides, LDL, leptin, and body length revealed significant associations with WGCNA modules, none of these overlapped with age (Figure 1).

**Figure 1.**
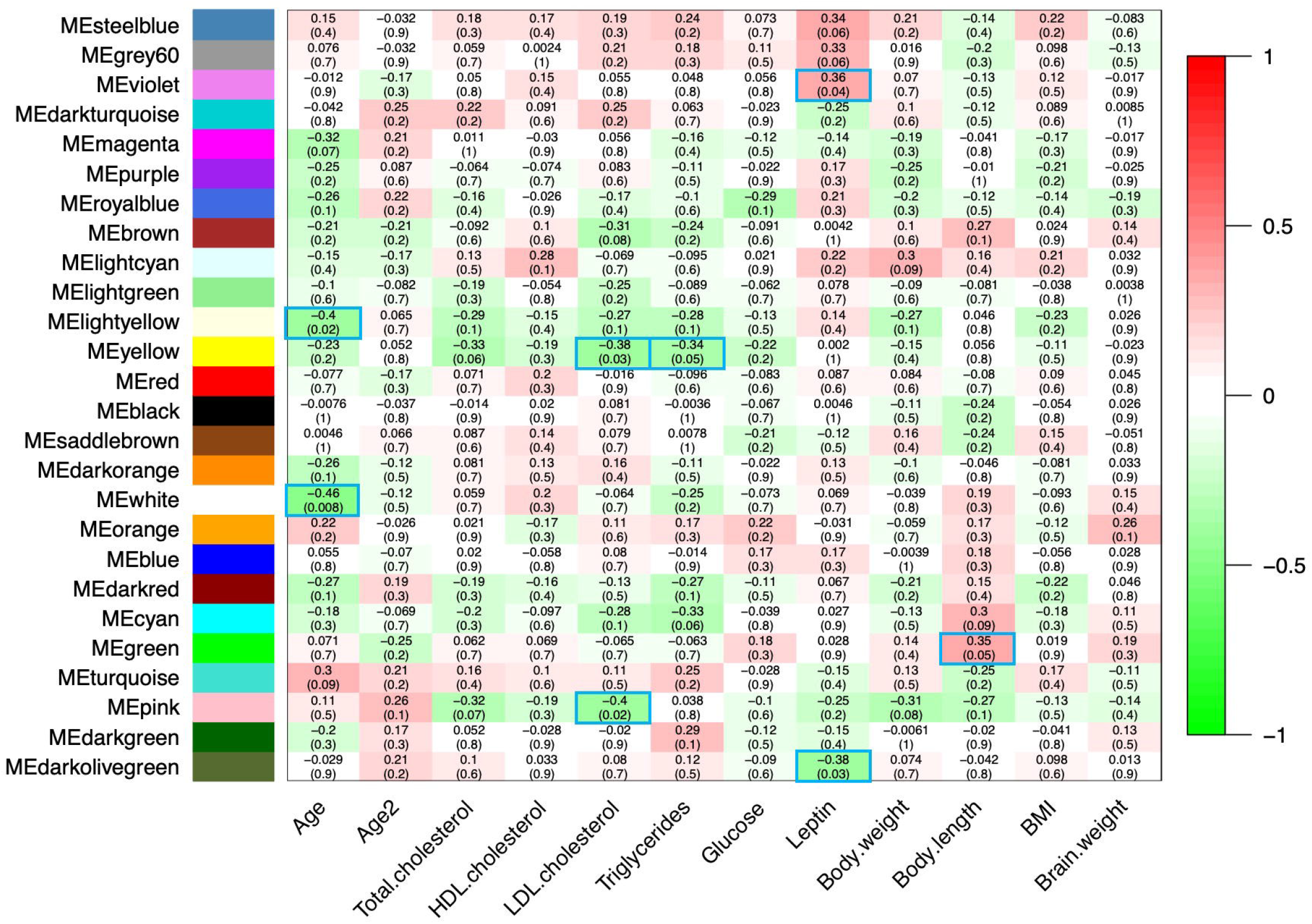
Modules associated with age and age-correlated traits. Each row corresponds to a module, the bottom column labels indicate each quantitative trait correlated with omic modules. Numbers in each box indicate correlations with p-values in parentheses. Positive correlations are indicated with red fill, negative with green fill, and significant modules are outlined blue.

### 3.3 Pathway Enrichment Analysis

Pathway analysis of genes, proteins, and metabolites correlated with age revealed 52 pathways with p-values < 0.01 and z-scores predicting directionality (Supplemental Table 8). Among these, the top 25 pathways all passed EOP filtering, all shared 5 or more genes in common, indicating extensive molecular coordination among the age-associated molecules in the PFC. Neurotransmitter signaling and other nervous system signaling pathways were the most statistically significant with the greatest numbers of molecules (Figure 2).

**Figure 2.**
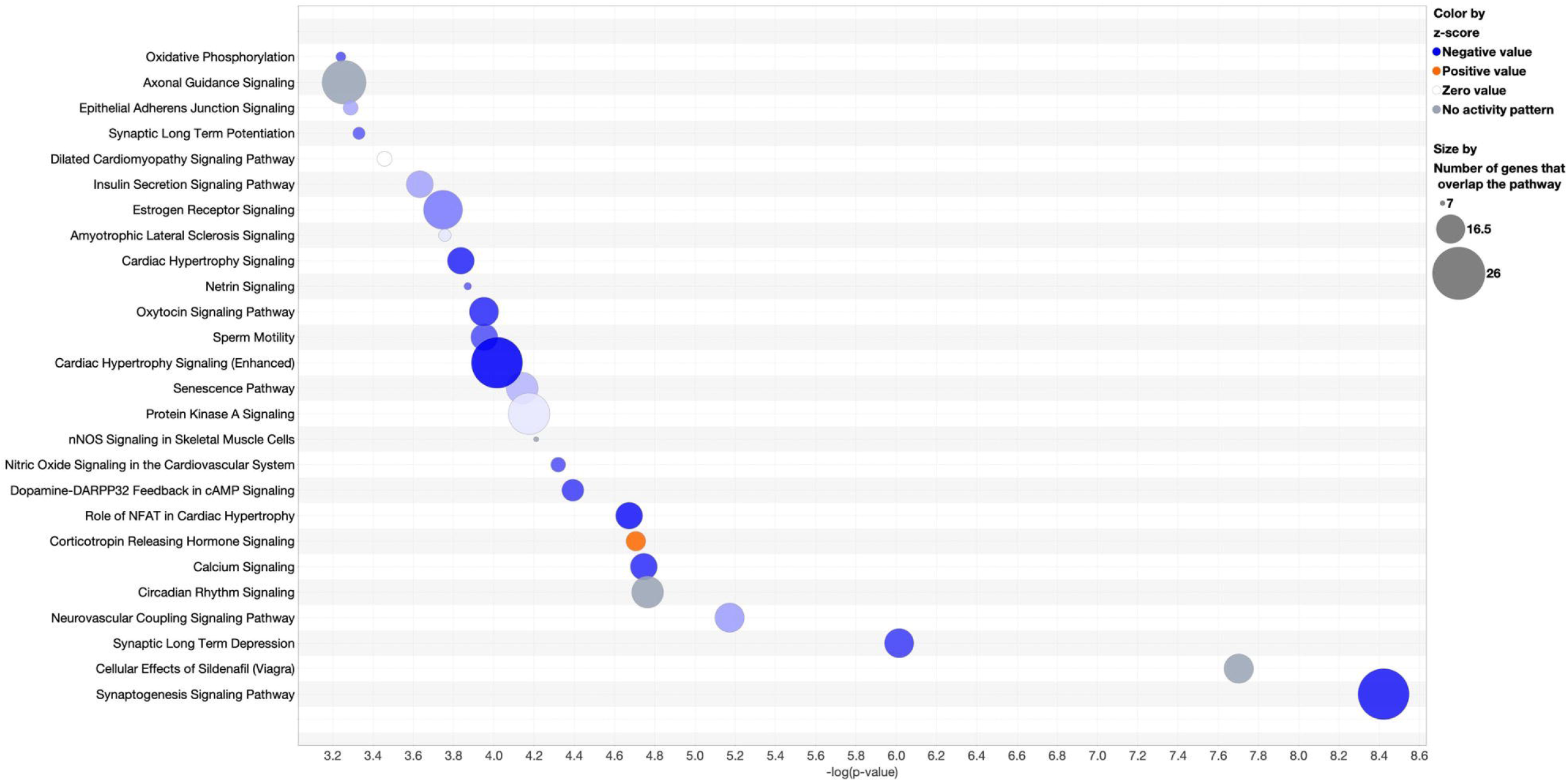
Bubble chart of significantly enriched, overlapping pathways with ≥ 5 genes in common. Pathways with enrichment -log (p-value) > 3.2 and absolute z-score >1.0 are shown. The x-axis shows the -log (p-value) and y-axis shows the pathways. Blue fill indicates pathways decreasing with age, orange indicates pathways increasing with age, and size of the circle denotes the number of omic molecules in the pathway.

The top 5 pathways highlighted our findings for signaling pathways previously associated with age in PFC, and novel pathways in PFC not previously associated with age. Consistent with previous studies (Kelly et al., 2014; Garcia et al., 2014), we found decreased abundance with age for *VEGFA* (vascular endothelial growth factor A) and *PDE1B* (phosphodiesterase 1B) genes and in dopamine signaling. In addition, two metabolites, γ-aminobutyric-acid (GABA) and 1,10-phenanthroline, in the dopamine signaling pathway were positively correlated with age (Figure 3), consistent with previous reports (Kiemes et al., 2021; Gao et al., 2013; Maitra et al., 2021). Also consistent with previous work was decreased activity of estrogen signaling with age (reviewed in (Friedman and Robbins 2022), including downregulation of IGF1 (insulin like growth factor 1), IGFR (insulin like growth factor 1 receptor), VEGFA, and EIF4EBP2 (eukaryotic translation initiation factor 4E binding protein 2) gene expression, and EIF4EBP1 (eukaryotic translation initiation factor 4E binding protein 1) and PRKAR1A protein expression (Supplemental Figure 1).

**Figure 3.**
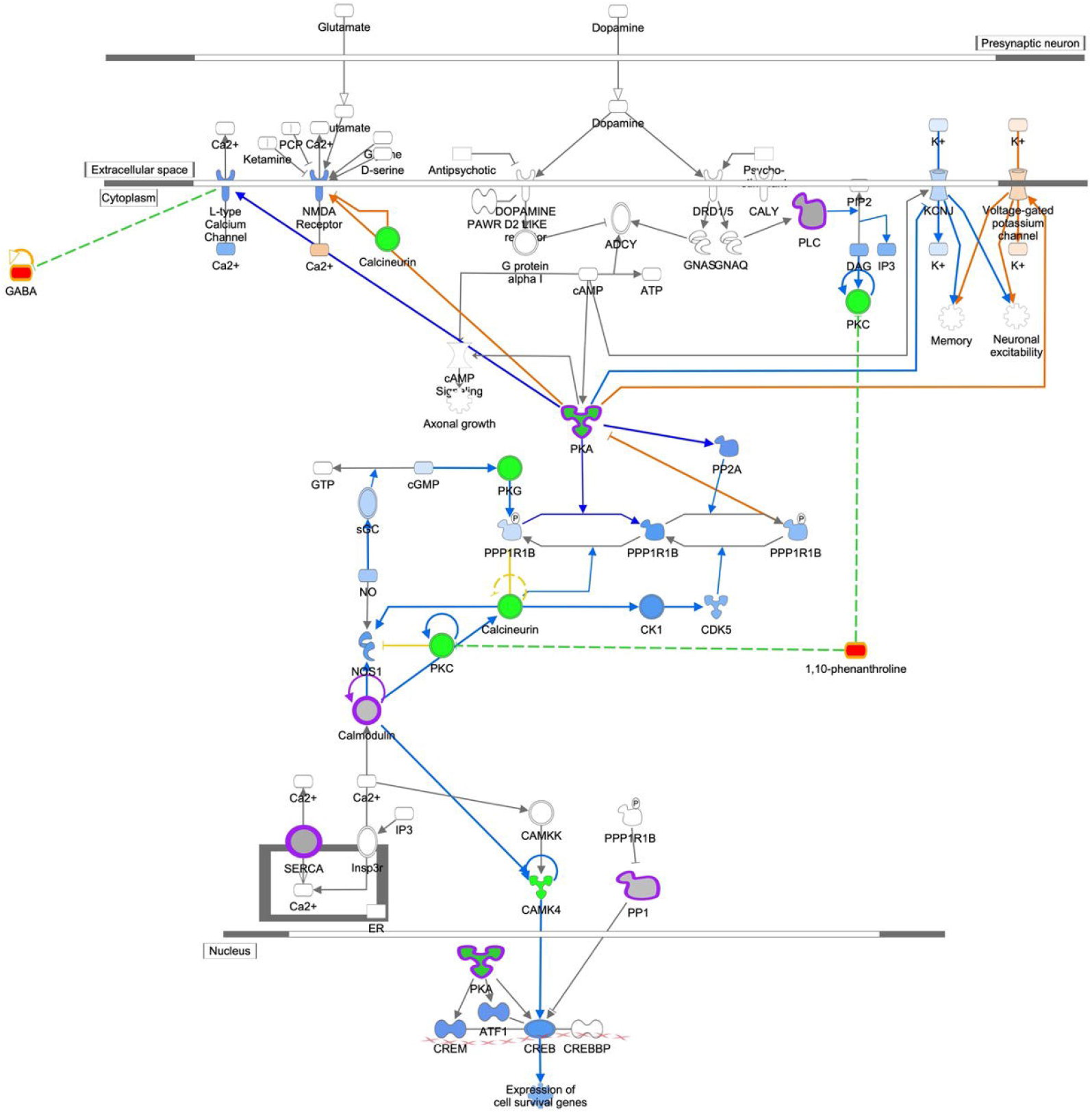
Dopamine DARPP32 Feedback in cAMP Signaling. Genes are indicated by green outline, proteins by purple outline, metabolites by gold outline, red fill indicates increased abundance with age, green fill decreased abundance with age, and gray fill indicates no change in abundance. Pathway enrichment p-value = 2.0^-06^.

PFC pathways associated with age included nitric oxide signaling (Figure 4), synaptogenesis (Supplemental Figure 2), and synaptic long-term depression (Supplemental Figure 3). The nitric oxide signaling pathway was significantly enhanced by including proteomic data, an example of proteomic data enriching the pathway beyond transcripts alone, with proteins found in key signaling steps such as calmodulin processing. In addition, multiple metabolites from the metabolomic dataset were associated with this pathway as regulators (1,10-phenanthroline and physostigmine) and by-products (GABA).

**Figure 4.**
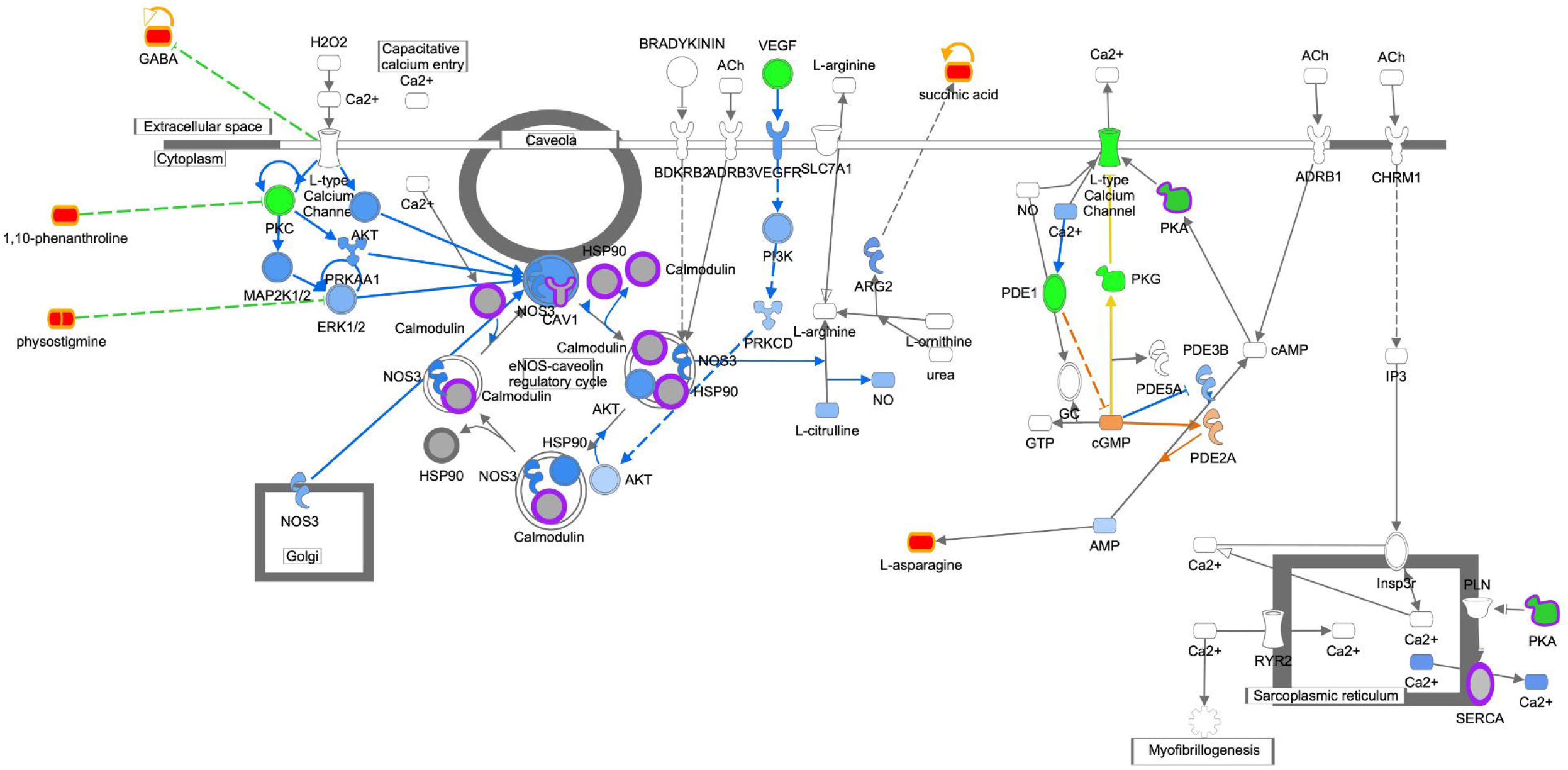
Nitric Oxide Signaling. Genes are indicated by green outline, proteins by purple outline, metabolites by gold outline, red fill indicates increased abundance with age, green fill decreased abundance with age, and gray fill indicates no change in abundance. Pathway enrichment p-value = 4.0^-06^.

Increased abundance of these metabolites in the PFC with age was inverse to decreased activity of the signaling pathway with age, consistent with previous studies (Kiemes et al., 2021; Gao et al., 2013; Maitra et al., 2021). Synaptogenesis signaling also included proteins in key signaling steps such as cadherin (CDH13) and EIF4EBP1. Identification of GABA and 1,10-phenanthroline inhibiting synaptic long-term depression signaling was also consistent with decreased PFC neuron activity with age in this cohort. Taken together, pathway enrichment analysis revealed coordinated decreased activity of neuronal signaling pathways with age in the primate PFC.

## 4 Discussion

The overall goal of this study was to use unbiased multi-omics analysis methods and data integration to identify molecular pathways associated with age in the female primate PFC. Previous studies have established the importance of the PFC in cognition (Miller and Cohen 2001) and shown commonalities in cognitive decline with age among humans and multiple NHP species (Lacreuse et al., 2020). Although ongoing studies have used RNA-Seq in human (Miller et al., 2017) and NHP (Wei et al., 2015; Li et al., 2013; Shao et al., 2010; Yuan et al., 2012; Somel et al., 2011; Li et al., 2019; Yin et al., 2020) brain region samples to quantify age-associated molecular changes, no studies to date have systematically characterized the broad adult age span in females included in this study, approximating a human age range of ∼30y to 88y. In addition, the size of our study cohort, which averages almost 3 animals per year of age, is far larger than any previous NHP study related to cognition and aging (reviewed in (Lacreuse et al., 2020; Upright and Baxter 2021). This study is unique in that animals were maintained on a low cholesterol, low fat “chow” diet throughout life, housed under identical conditions, and neural tissues were collected on a defined schedule with short PMIs (∼30 min) which allowed us to assess PFC molecular changes associated with normal aging across multiple omics domains.

We quantified morphometry and clinical chemistries for measures previously positively associated with age-related co-morbidities and found measures of body length associated with age, as well as plasma LDL, triglyceride concentrations, and glucose. Interestingly, we did not find significant associations between either HDL or total serum cholesterol and age in this cohort of female baboons.

Using unbiased WGCNA, we identified 2 modules of omics molecules that were significantly correlated with age. Neither of these age-associated modules overlapped with triglycerides, or morphometric measures correlated with age, suggesting that molecular pathways influencing age-associated variation in these morphometric and clinical measures differ from molecular pathways influencing PFC age-associated changes. Overall, the number of molecules in the two age-correlated WGCNA modules was small, including only 2.3% of transcripts and 1.4% of proteins that passed quality filters. Although few metabolites correlated with age, 5/20 (25%) were remarkably informative and consistent with gene/protein pathway directionality. Furthermore, pathway enrichment analysis showed extensive overlap and coordination based on molecules common among the top 25 pathways and consistent with decreased age-associated activity. Taken together, these results provide a detailed molecular picture in which few molecules were associated with age, but those few were highly interconnected, suggesting coordinated functional changes with age in the primate PFC.

Our unbiased integrated omics approach identified both known and novel pathways associated with PFC aging in primates, and provides evidence for molecular mechanisms mediating age-related cognitive decline and reduced PFC function. Known pathways include PFC dopaminergic circuits, which are essential for cortical connectivity and cognition (Arnsten et al., 2012; Dehaene and Changeux 2000), and have been shown in imaging studies to decrease with age, consistent with decreased dopamine signaling reported here (Berry et al., 2016). GABA-mediated inhibition is critical to the function of neural circuits that support memory in the PFC (Deco and Rolls 2003; McQuail et al., 2015). Here, GABA was positively correlated with age. GABA and dopamine signaling act together to support spatial working memory in the primate dorsolateral PFC (Arnsten et al., 2015; Goldman-Rakic 1995). Also, GABA is negatively correlated with human neural activity in functional MRI (Kiemes et al., 2021; Gao et al., 2013) which is consistent with our findings of increased GABA abundance with increased age. We also observed decreased estrogen signaling with age, which has been associated with cognitive decline (reviewed in (Friedman and Robbins 2022) and modulates dopamine (Shanmugan and Epperson 2014) and GABA (Gilfarb and Leuner 2022) function. Proteins and metabolites in these pathways were of particular interest, providing multi-level molecular data on key signaling steps and pathway read outs, respectively.

The top ranked pathways for primate PFC included decreased nitric oxide signaling with age. Our results differ from a study in male rats showing increased eNOS with age (Liu et al., 2004). It is possible this difference is based on significant differences in PFC cell composition and function between rodents and primates (Seamans et al., 2008) or may be due to sex-specific signaling, i.e., female NHP versus male rodents. Indeed, estrogen induces synaptogenesis in the primate PFC (Hara et al., 2015). GABA and 1,10-phenanthroline may contribute to synaptic long-term depression by inhibition of glutamate at the synaptic vesicle and protein kinase C, respectively, key molecules in the signaling pathway. Not only is GABA negatively correlated with neural activity, 1,10-phenanthroline has been shown to promote apoptosis (Maitra et al., 2021). Increased abundance in these metabolites with age are consistent with decreased signaling activity in the synaptic long-term depression signaling pathway. Decreased signaling with age in these pathways combined with decreased dopamine, GABA, and estrogen signaling indicate decreased PFC neuronal activity, even in healthy primate aging, which are likely involved in common declines in cognitive function observed with aging.

Also noteworthy is that previous human studies of molecular changes in PFC with age have reported changes related to inflammation and immune function (Zhang and Wong 2022; Tennakoon et al., 2022; Wruck and Adjaye 2020), and reported evidence of age-associated differences in immune function in PFC. However, the analyzed tissue samples were derived from seven studies with high heterogeneity in patient populations, cause of death, and PMI. We did not find these pathways enriched in our dataset which was derived from NHPs that consumed a healthy, low-fat, low-cholesterol, largely plant-based diet throughout life, and in which PFC was collected rapidly, using the same protocol for all subjects. Thus, it is difficult to untangle lifetime diet effects, the use of tissues from healthy subjects, and PMI as factors accounting for these differences.

### 4.1 Limitations

Although this study is the first of its kind for cohort size, age span, and integrated omics, there are limitations that should be addressed in future work. First, this study included only females, due to the difficulty of maintaining large numbers of males in NHP colonies, as non-breeders are typically culled prior to aging. Thus, the extent to which the observations reported here are sex-specific remains to be determined. Second, all omics methods were “bulk” methods for this heterogeneous brain region – we likely were not able to identify all pathways associated with age as molecular signaling in less abundant cell types would likely be undetected. We also did not characterize in more detail post-translational modifications in proteins, a future analysis that may be of interest given the key role of several proteins in the identified signaling pathways where alterations in phosphorylation may further impact activity of key protein regulators identified in our study. Finally, although reproductive status was not characterized, previous studies show that baboon females may become peri- or post-menopausal as early as 20 years of age, consistent with the decline in estrogen signaling with age (Macrini et al., 2013).

## 5. Conclusions

Nonetheless, our unbiased integrated omics analysis of the female primate PFC revealed novel neural signaling pathways in which activities decrease with age. Analysis of PFC samples collected from animals maintained on a healthy diet in social groups throughout lifespan provides a framework for normal healthy aging in the PFC at the molecular level. As mentioned previously, integration of metabolomic data provides useful inputs and readouts from signaling pathways. In addition, because GABA can be quantified by current imaging modalities (Kiemes et al., 2021; Gao et al., 2013), the association of GABA with these known and novel age-associated signaling pathways provides one potential biomarker to assess PFC changes with age and in response to stressors. These signaling pathways, and their changes with aging, are likely to represent critical molecular contributors to age-related cognitive decline and overall PFC function, even prior to overt clinical symptoms, and as such may provide novel insights into age-related disease processes. Future work is required to identify master regulators that mediate the decreased activity of these highly interconnected neuronal signaling pathways, and to identify sex differences.

## 7 Declaration of generative AI in scientific writing

AI and AI-assisted technologies were not used in the writing process.

## 8 Author Contributions

LAC, GDC, PWN, and MO contributed to conception and design of the study. SP, JC, ZH, IA, HFH, GL, AYLJ, BW, TCR, and CL, generated and analyzed omic data. KDZ, and SP performed statistical analyses. LAC wrote the first overall draft of the manuscript. All authors contributed to manuscript revision, and read and approved the submitted version.

## 10 Acknowledgements

The authors would like to also acknowledge the staff at the Southwest National Primate Research Center, Texas Biomedical Research Institute for the veterinary services and animal maintenance support.

## 13 Supplementary Material

### 13.1 Supplemental Figures

**Supplemental Figure 1.**
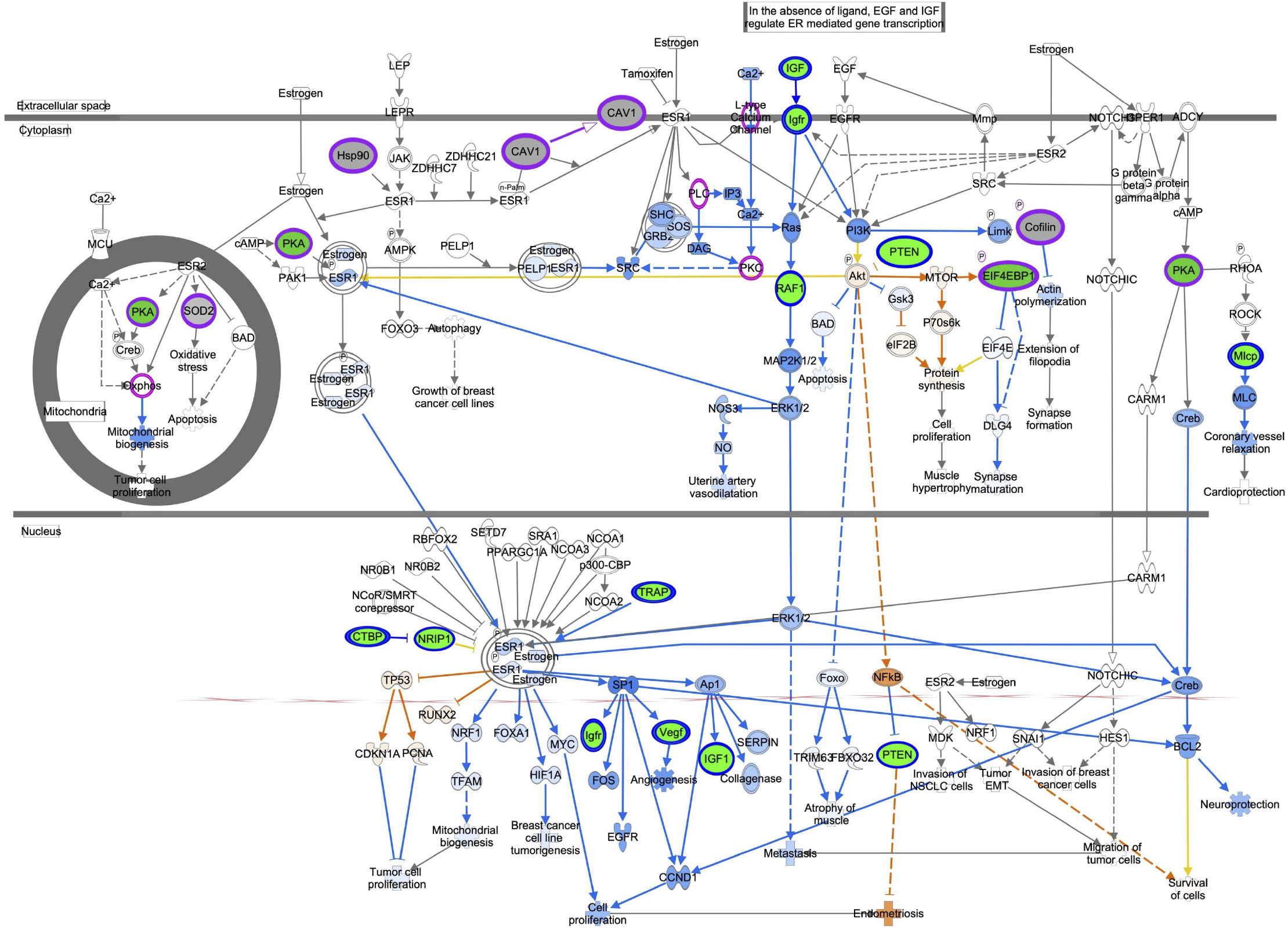
Estrogen Receptor Signaling. Genes are indicated by blue outline, proteins by purple outline, metabolites by gold outline, red fill indicates increased abundance with age, green fill decreased abundance with age, and gray fill indicates no change in abundance. Pathway enrichment p-value = 7.9^-06^.

**Supplemental Figure 2.**
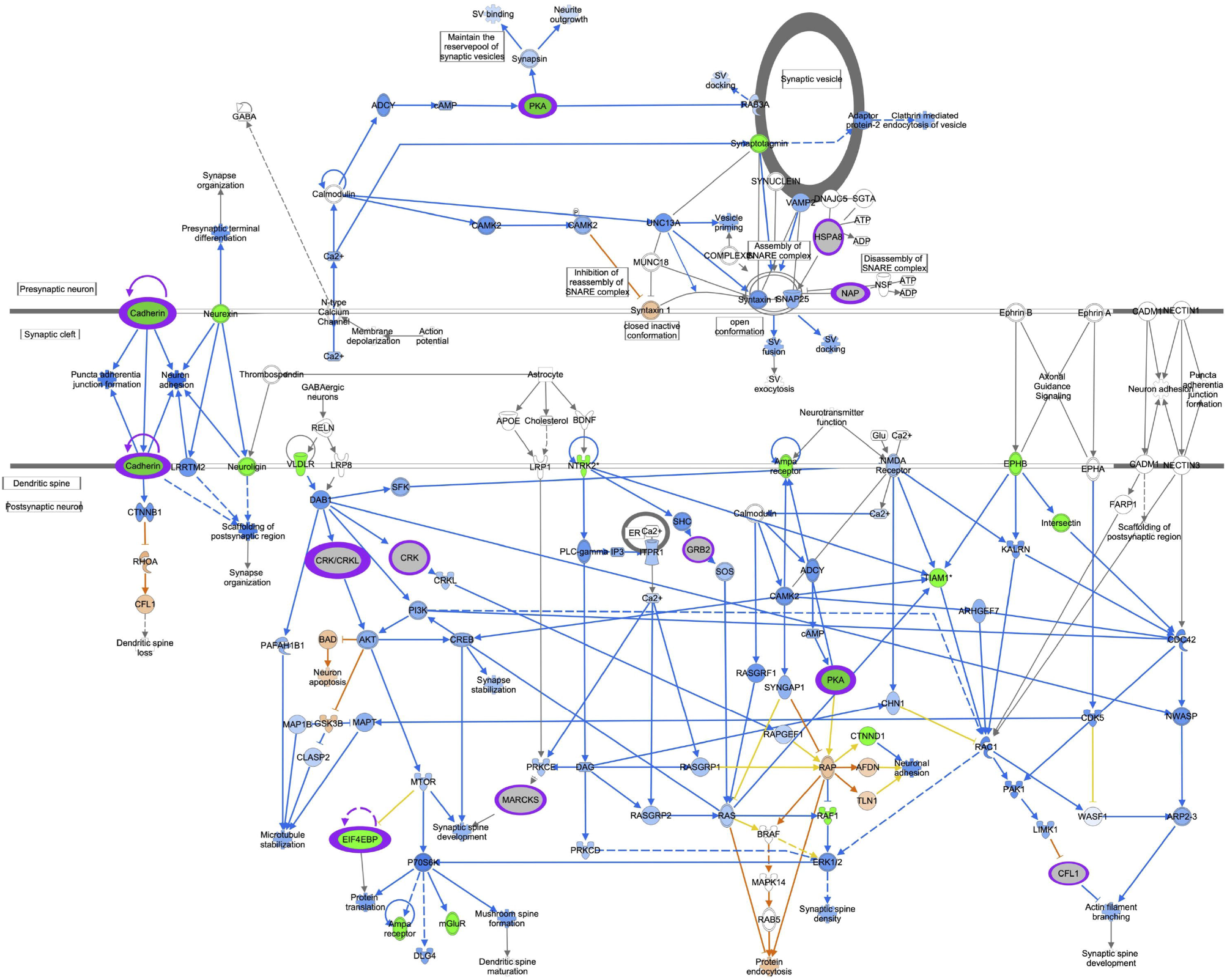
Synaptogenesis Signaling. Genes are indicated by blue outline, proteins by purple outline, metabolites by gold outline, red fill indicates increased abundance with age, green fill decreased abundance with age, and gray fill indicates no change in abundance. Pathway enrichment p-value =7.9^-08^.

**Supplemental Figure 3.**
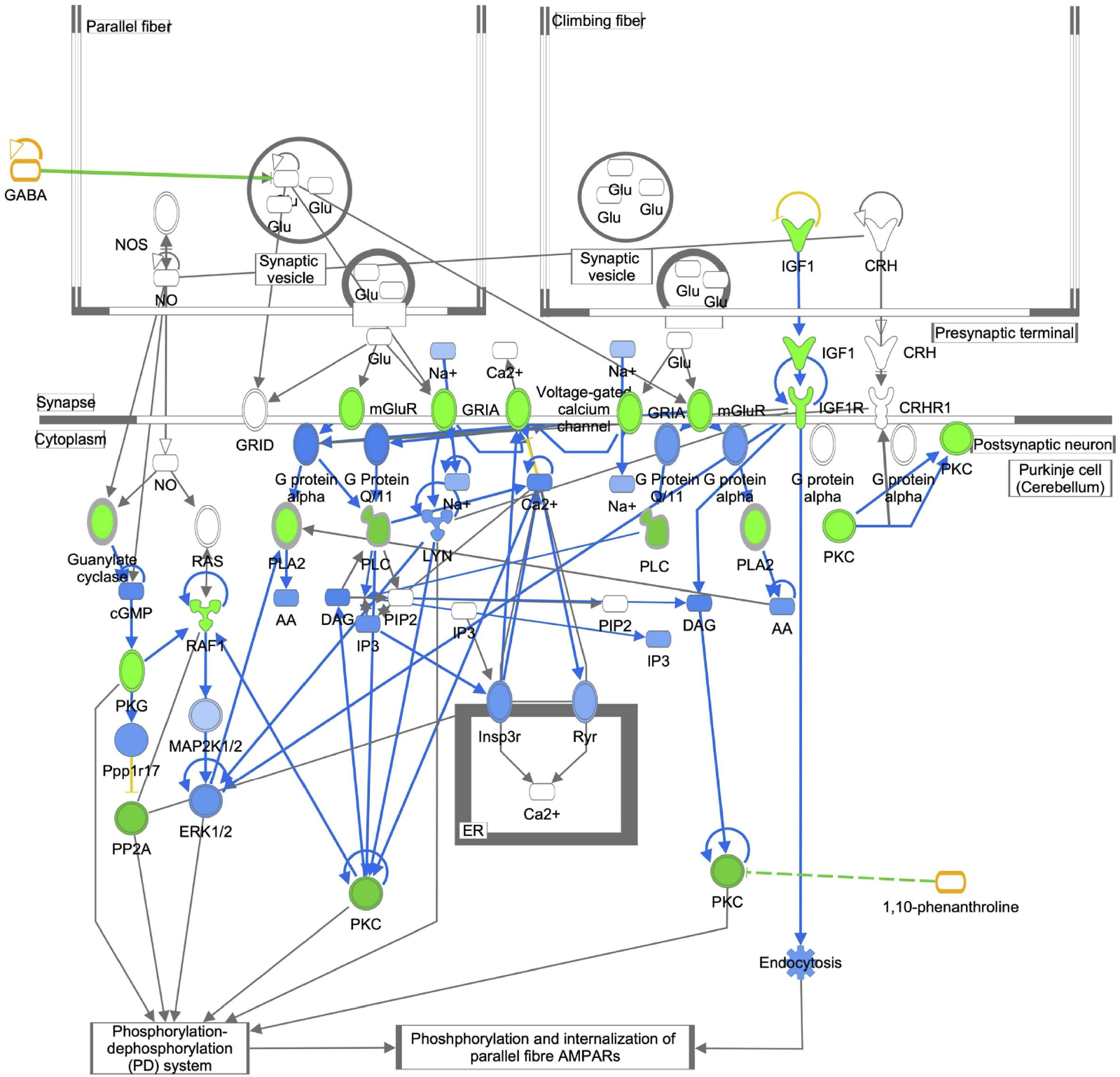
Synaptic Long -Term Depression Signaling. Genes are indicated by blue outline, proteins by purple outline, metabolites by gold outline, red fill indicates increased abundance with age, green fill decreased abundance with age, and gray fill indicates no change in abundance. Pathway enrichment p-value =1.3^-08^.

### 13.2 Supplemental Tables

**Supplemental Table 1**. Clinical and morphometric measures.

**Supplemental Table 2**. Normalized gene expression.

**Supplemental Table 3**. Normalized protein abundance.

**Supplemental Table 4**. Normalized metabolite abundance.

**Supplemental Table 5**. WGCNA modules with significantly correlated phenotypic measures.

**Supplemental Table 6**. Genes, proteins, and metabolites in the age associated white and lightyellow modules.

**Supplemental Table 7**. Regression analyses for age-protein, age squared-protein, age-metabolite, and age squared-metabolite.

**Supplemental Table 8**. Pathway enrichment analysis.

